# The Genomic Legacy of Ancient Polyploidy in Crop Domestication

**DOI:** 10.64898/2026.03.09.710542

**Authors:** Michael McKibben, Michael Barker

## Abstract

Species that have an ancestry of whole-genome duplications (WGDs) are more likely to be domesticated, but the underlying mechanisms remain unclear. We tested whether paleologs—genes duplicated during ancient WGDs—are enriched in candidate domestication lists across 22 crop species spanning 17 genera. Paleologs were significantly enriched in 14 species, with single-copy paleologs showing the most consistent overrepresentation. In contrast, genes from small-scale duplications were consistently underrepresented among domestication candidates. This enrichment was independent of WGD age and degree of gene loss, indicating that constraint on copy number does not preclude selection on gene function. Several non-mutually exclusive processes could explain this pattern, including accumulated genetic diversity becoming available upon return to single-copy, selection to maintain essential functions, and greater selection efficiency on unmasked loci. Ancient WGDs thus provide a persistent genomic substrate for crop evolution millions of years later.

## Introduction

Polyploidy, or whole genome duplication (WGD), is a fundamental evolutionary force among plants and has significantly impacted agricultural development. Polyploid plants are more than twice as likely to be domesticated than their diploid relatives.^1^ This relationship is thought to arise from increased morphological variation and genomic plasticity found in polyploids, including enhanced heterozygosity, altered recombination landscapes, and novel gene expression patterns.^2–6^ These features may explain why polyploids exhibit higher rates of niche differentiation and adaptation to environmental stress compared to their diploid counterparts.^7–9^ However, less is known about the impact of paleopolyploid ancestry on crop domestication. All diploid crops have a polyploid ancestry, having undergone an average of approximately 3.5 WGD events throughout their evolutionary history.^10,11^ Genomic signatures of these ancient events persist for millions of years.^11–13^ However, whether the advantageous features of polyploidy for domestication persist following diploidization remains an open question.

During diploidization, alleles on different chromosomes within a polyploid genome can be converted to separate loci. These duplicated genes—termed ohnologs or paleologs—can persist for millions of years.^13–15^ and have several potential consequences for crop domestication. Initially, duplicate copies maintain dosage balance with other duplicated genes in the genome.^16–19^ This redundancy can mask loci from selection, mitigating the negative impacts that gene deletions and deleterious alleles can have on yield.^20,21^ Similarly, masking can enable duplicate loci to accumulate genetic diversity that may be advantageous during domestication.^22^ While this elevated genetic diversity is generally only exposed when duplicates return to the single copy state, instances of sub- and neofunctionalization demonstrate that this variation can also be exposed through functional diversification.^22–27^ However, masking is not a ubiquitous feature of duplicate loci, as some gene functional categories experience purifying selection for absolute dosage.^19,28–30^ These processes are not mutually exclusive, and their relative contributions to crop domestication likely depends on species-specific selective contexts. For example, duplicate genes contribute to repeated global ^31^, but not local ^32^, adaptation to abiotic stress. Crop domestication may be a similar situation where several common traits, called domestication-syndromes, are selected across numerous species.^33–35^

Recent work in *Brassica rapa* revealed that paleologs contain elevated levels of genetic diversity relative to other genes and are over-represented in candidate domestication gene lists made using several methods.^22^ Both multi-copy and single-copy paleologs were overrepresented among differentially expressed genes between cultivars and genes detected under selective sweeps.^22^ However, candidate genes that were detected using direct tests of selection (McDonald–Kreitman) were only enriched with single-copy paleologs.^22^ This pattern may be a result of methodological differences, or an untested feature of copy-number variation that impacts how loci respond to selection. Furthermore, it remains unknown whether paleologs are unique or if other duplicate-gene classes also contribute significantly to domestication. The predominant mode of duplication likely depends on the underlying protein-protein interactions (PPI) and gene-regulatory networks (GRN) that determine traits of interest for artificial selection.^36,37^ Each form of duplication is known to influence the topology of these networks differently^37,38^, making it difficult to *a priori* determine if the results found in *B. rapa* can be extended to other species and traits of interest. Determining whether these patterns can be generalized across angiosperms requires examining broader analyses of how genes evolve following WGD.

The contribution of paleologs to domestication may depend on the selective constraints acting on them following WGD. A large class of ‘core’ angiosperm genes consistently returns to single-copy status following duplication, a pattern attributed to strong purifying selection against copy number changes due to dosage sensitivity.^39^ These rapidly-returning genes show slower rates of sequence and expression divergence, leading to suggestions that dosage constraints may limit the ease with which genome multiplications can facilitate evolutionary innovation.^40^ The dosage balance hypothesis posits that purifying selection can limit functional divergence between duplicate gene copies^19^, and recent reviews have noted that dosage balance requirements may constrain the potential of whole genome duplication to promote evolutionary change.^41^ Based on these observations linking rapid return to single-copy status with constrained evolution, genes under strong dosage constraint have been considered unlikely targets for the positive selection driving adaptation and domestication. Whether this inference holds in a domestication context has not been tested.

Here, we tested if candidate genes associated with domestication show overrepresentation of paleologs or other duplicate-gene classes across multiple crop species. We identified 22 agronomically important species across 17 genera (Table 1) with published candidate gene lists from population genomic and quantitative genetic analyses. We classified genes by first identifying paleologs using the machine learning approach Frackify.^42^ Non-paleologs were then classified as singletons, tandem, segmental, and proximal duplicates using MCScanX.^43^ Using these classifications, we assessed whether paleologs or other duplicate-gene classes were enriched among candidate domestication genes. Dosage-balance is often evoked when predicting how paralogs respond to selection, a conclusion motivated in part by observed biased retention of specific Gene-Ontology (GO) categories among paralog classes.^19,28–30,44–46^ We leveraged GO terms to assess if the distribution of duplicate-gene classes amongst each genome and candidate list are consistent with dosage-based predictions. For six species from our analysis that overlapped with the Li et al.^39^ and Tasdighian et al.^40^ analyses we assessed if paralog duplicability influenced their propensity to be over-represented in candidate gene lists. Finally, given the age of WGDs in our species varied greatly, we assessed if time since duplication or degree of gene loss correlated with the over- or underrepresentation of paleologs in our candidate domestication gene lists. Our findings suggest that paleologs, particularly single-copy paleologs, represent the most consistently overrepresented gene-duplication class among candidate domestication genes regardless of age.

## Results

### Literature survey of candidate domestication gene lists

We surveyed the literature using Google Scholar and identified 21 studies representing 22 species across 17 genera (Figure 1, Table 1) with publicly available candidate domestication gene lists that matched associated reference genomes. Candidate genes were inferred in these studies by a variety of approaches including Reduction of Diversity (ROD) metrics such as π, or Quantitative Trait Loci (QTL) and Genome Wide Association studies (GWAS) comparing landrace to domesticated individuals (Table 1). More than one approach was used to identify candidate genes in 15 of the 21 studies. Across our survey, candidate gene lists ranged from 55 (*Brassica juncea*) to 5,605 genes (*Solanum lycopersicum*). In total, we collected 30,928 candidate crop domestication and improvement genes from the literature.

**Figure 1:**
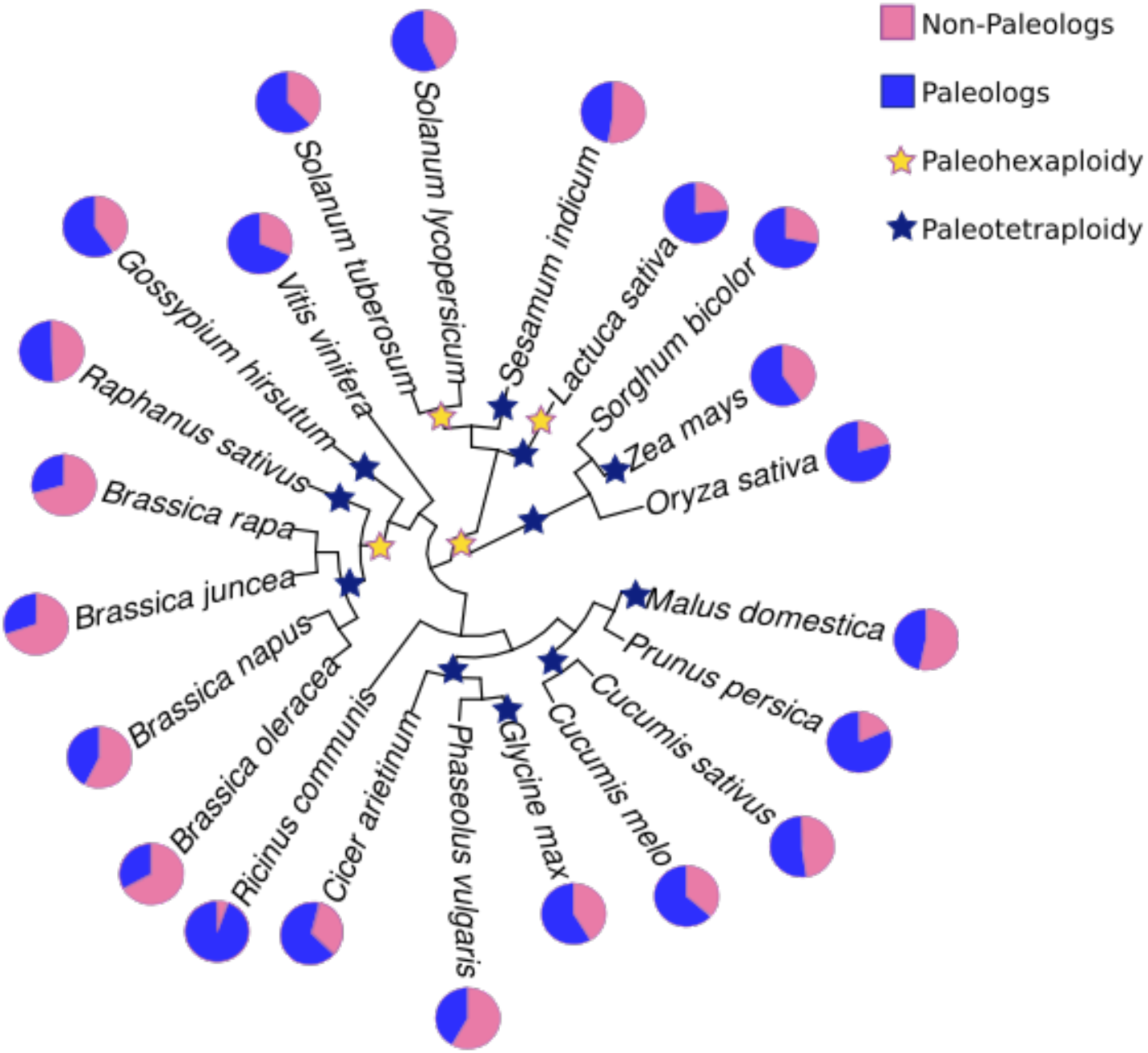
The basic phylogeny of species used in this analysis.^76^ Pie charts represent the ratio of paleologs to non-paleologs from the most recent WGD in each species. Stars denote the approximate placement paleotetraploid and paleohexaploid WGD based on 1KP analyses.^10^

### Identification of genomic regions originating from ancient WGD

We identified paleologs originating from the most recent WGD in each species using the Frackify pipeline.^42^ Approximately 50.2% (462,657) of the 921,086 annotated genes across all our species were classified as paleologs originating from the most recent WGD in their respective species (Figure 2, Supplementary Table 2-3). Paleolog retention varied substantially, from 22.9% of genes in *Oryza sativa to* 70.5% in *B. rapa,* averaging 44.4% of genes across all species (Figure 2, Supplementary Table 2-3). Members of Brassicaceae retained the highest fraction of their genome as paleologs, averaging 60.2% across three diploid species tested. We also identified paleologs from ancient WGD in four polyploid species, *Brassica juncea, Brassica napus, Gossypium hirsutum,* and *Glycine max* (Figure 3, Supplementary Table 2-3), which retained on average 63.4% of their genomes as paleologs.

**Figure 2:**
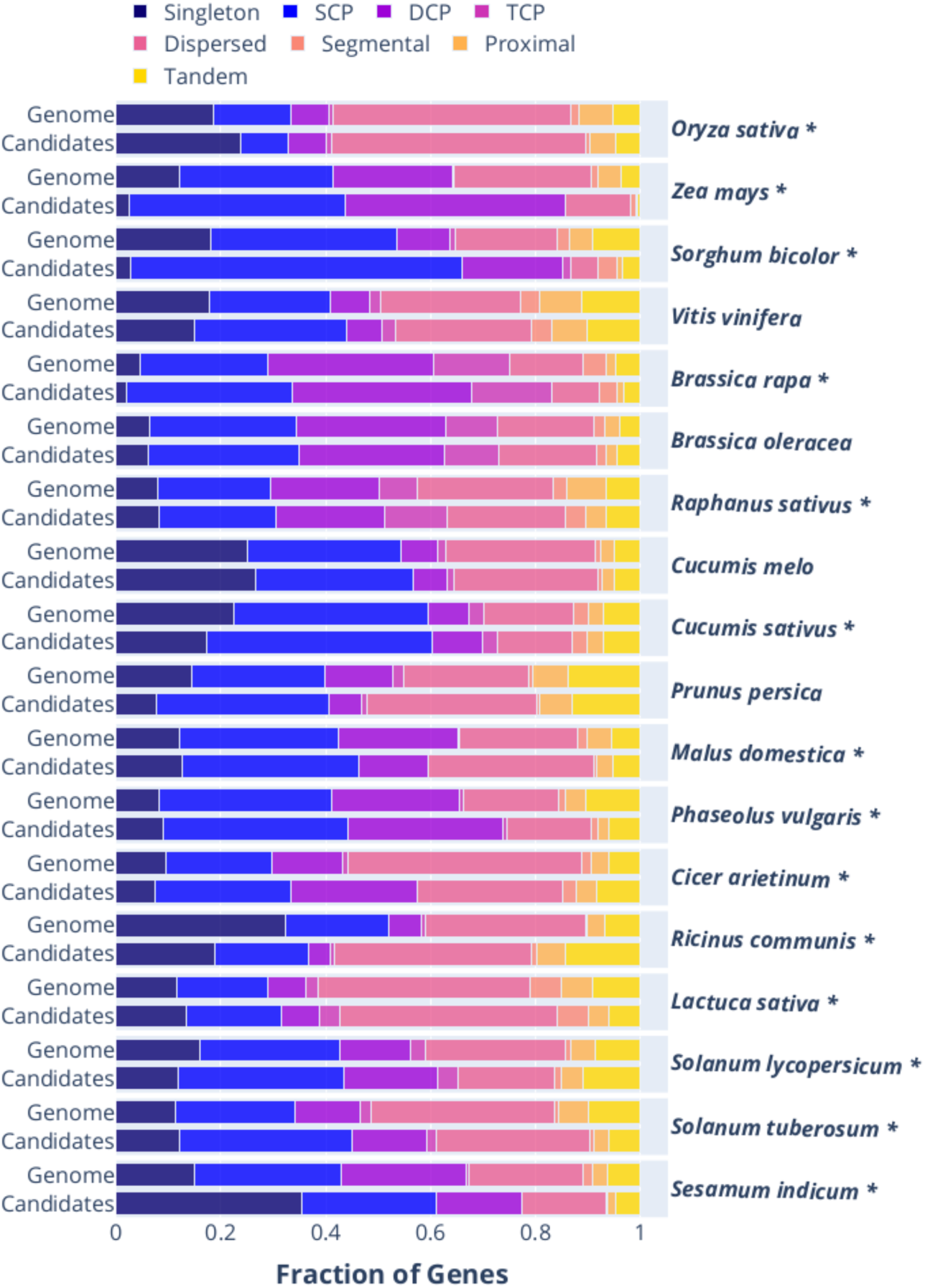
Bar charts depicting the distribution of duplicate-gene classes across the genomes of diploid species and their associated candidate domestication gene lists. Species with asterisks were significant in a Pearson Chi-squared tests of goodness of fit comparing the distribution of duplicate-gene classes between each gene list.

**Figure 3:**
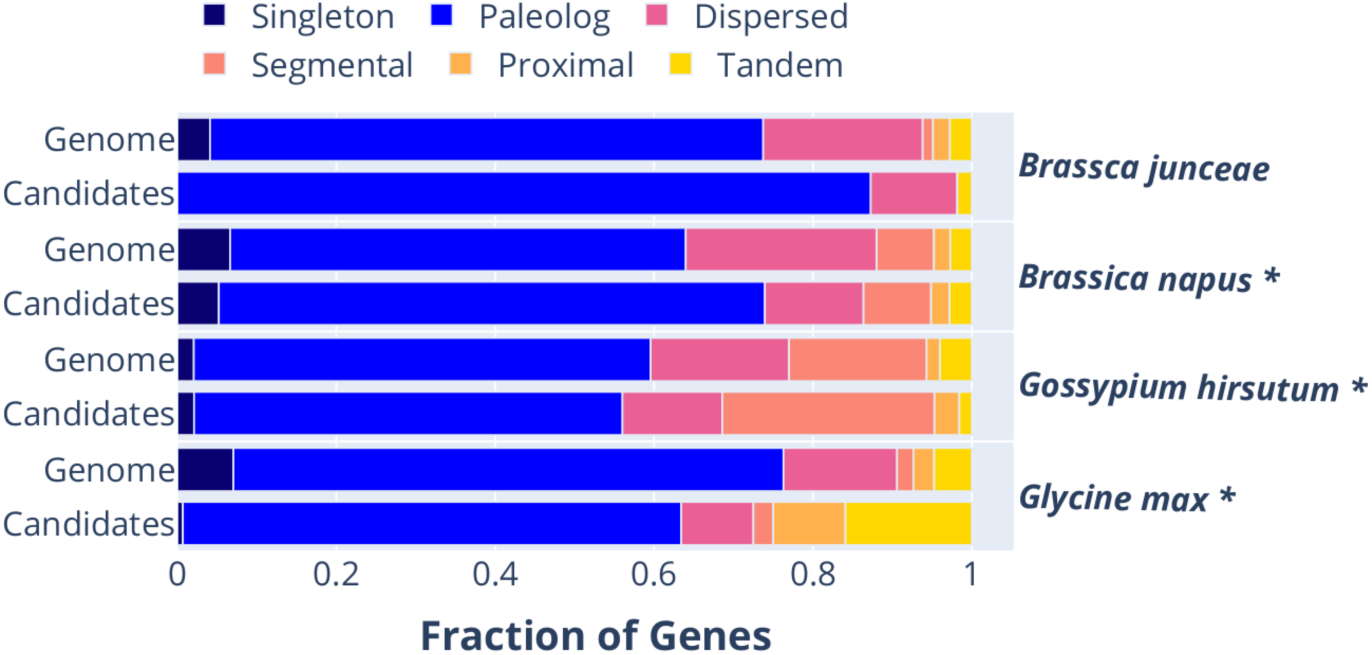
Bar charts depicting the distribution of duplicate-gene classes across the genomes of polyploid species and their associated candidate domestication gene lists. Species with asterisks tested significantly in Pearson Chi-squared tests of goodness of fit comparing the distribution of duplicate-gene classes between each gene list.

Following duplication, one or more copies of a paleolog may be lost through fractionation, resulting in a variety of retention statuses for duplicate genes. Frackify is able to classify paleologs by how many retained paralogous loci are present, be that all three paralogous loci (triple-copy) in hexaploids, two loci (double-copy), or only one surviving loci (single-copy). Across diploids, the majority of paleologs were retained in single-copy state (60.7% average), ranging between 34.5% in *B. rapa* to 77.9% *Zea mays* (Figure 2, Supplementary Table 2-3). The fraction of paleologs retained in duplicate had a narrower range compared to single-copy paleologs; varying from 16.4% in *Cucumis sativus* to 45.5% in *Sesamum indicum* (33.3% average)(Figure 2, Supplementary Table 2-3). The primary driver of increased paleolog retention in the Brassicaceae were triple-copy paleologs, averaging 16.7% compared to the average of 6.0% among Asteraceae, Cucurbitaceae, Euphorbiaceae, Solanaceae and Vitaceae. We did not assess the retention class of paleologs in the four polyploid species, as nested duplication histories makes it difficult to ascertain the precise retention patterns in their ancient WGDs.

### Identification of duplicate genes originating from small scale duplications

Gene families also expand by different kinds of small scale duplications (SSD) in addition to WGD. We used MCScanX to further classify the 458,429 genes Frackify classified as non-paleologs. MCScanX identified 355,488 (75.5%) of these non-paleologs as SSD (Figure 2, Supplementary Table 2-3), whereas the remaining 102,941 genes were classified as singletons. Across species, the majority of SSD were classified as dispersed duplicates, ranging from 52.9% in *Prunus persica* to 80.0% in *Cicer arietinum* (64.8% average)(Figure 2, Supplementary Table 2-3). On average, tandem duplicates made up 18.5% of SSD across species, with a maximum of 30.7% observed in *Phaseolus vulgaris* and minimum of 8.9% in *O. sativa* (Figure 2, Supplementary Table 2-3). There were relatively few proximal and segmental duplicates among SSDs, averaging 11.0% and 5.5% respectively (Figure 2, Supplementary Table 2-3). The Brassicaceae species *B. rapa* and *Raphanus sativus* had the highest fraction of segmental (17.4%) and proximal (18.1%) duplicates. In contrast, the lowest fractions of segmental and proximal SSD were observed in *Ricinus communis* (<1%) and *C. arietinum* (5.9%)(Figure 2, Supplementary Table 2-3).

### The distribution of gene duplication classes in candidate gene lists

To assess which gene-duplication classes played an important role during the domestication process, we used a Pearson chi-square test of goodness of fit between the distribution of duplicate-gene classes in candidate gene lists to the distribution in the entire genome. Among 18 diploid species tested, 15 had significantly different distributions of gene duplication classes between their genome and candidate lists (Figure 2, Supplementary Table 2,4). *Post hoc* testing on the residuals of each gene duplication class found that paleologs as a broad class were the most commonly overrepresented class, having a significant p-value in ten species (Figure 2, Supplementary Table 2,4). In our initial comparison, both single- and double-copy paleologs were significantly enriched in six out of the 18 diploids respectively. We observed the opposite pattern in SSD. Tandem and proximal duplicates were significantly underrepresented in six and five species respectively. Dispersed duplicates were underrepresented in seven species (Figure 2, Supplementary Table 2,4). Singleton genes followed a similar pattern to SSD, as six species had an under- and one (*S. indicum*) an overrepresentation of singletons in their candidate lists (Supplementary Table 2,4). These broad patterns were not universal, as three species had an underrepresentation of paleologs (*O. sativa, Malus domestica, and S. indicum*) and three species had an overrepresentation of SSD in their candidate gene lists (*M. domestica, R. communis, and Solanum lycopersicum*)(Figure 2, Supplementary Table 2,4). In four polyploid species tested, SSD were overrepresented in *Glycine max* and *Gossypium hirsutum*, and underrepresented in *Brassica napus* (Figure 3, Supplementary Table 3,5). *Brassica napus* and *G. max* also showed an over- and underrepresentation of paleologs, respectively (Figure 3, Supplementary Table 3,5).

To better understand which gene duplication classes were frequently over- and underrepresented across our entire dataset, we used a Friedman test on the ranked residuals from the species-specific chi-squared tests. Following a significant result (χ^2^ =20.61, *p*=0.004), we conducted a Nemenyi post-hoc test on each pairwise comparison of duplicate-gene classes. The majority of comparisons were non-significant with the exception of single-copy paleologs, which contributed to model significance more frequently than singletons, dispersed, and proximal duplicates (Figure 4, Supplementary Table 6). These results in combination with post-hoc tests for each chi square suggest that single-copy paleologs are the only gene-class to be consistently overrepresented in candidate gene lists.

**Figure 4:**
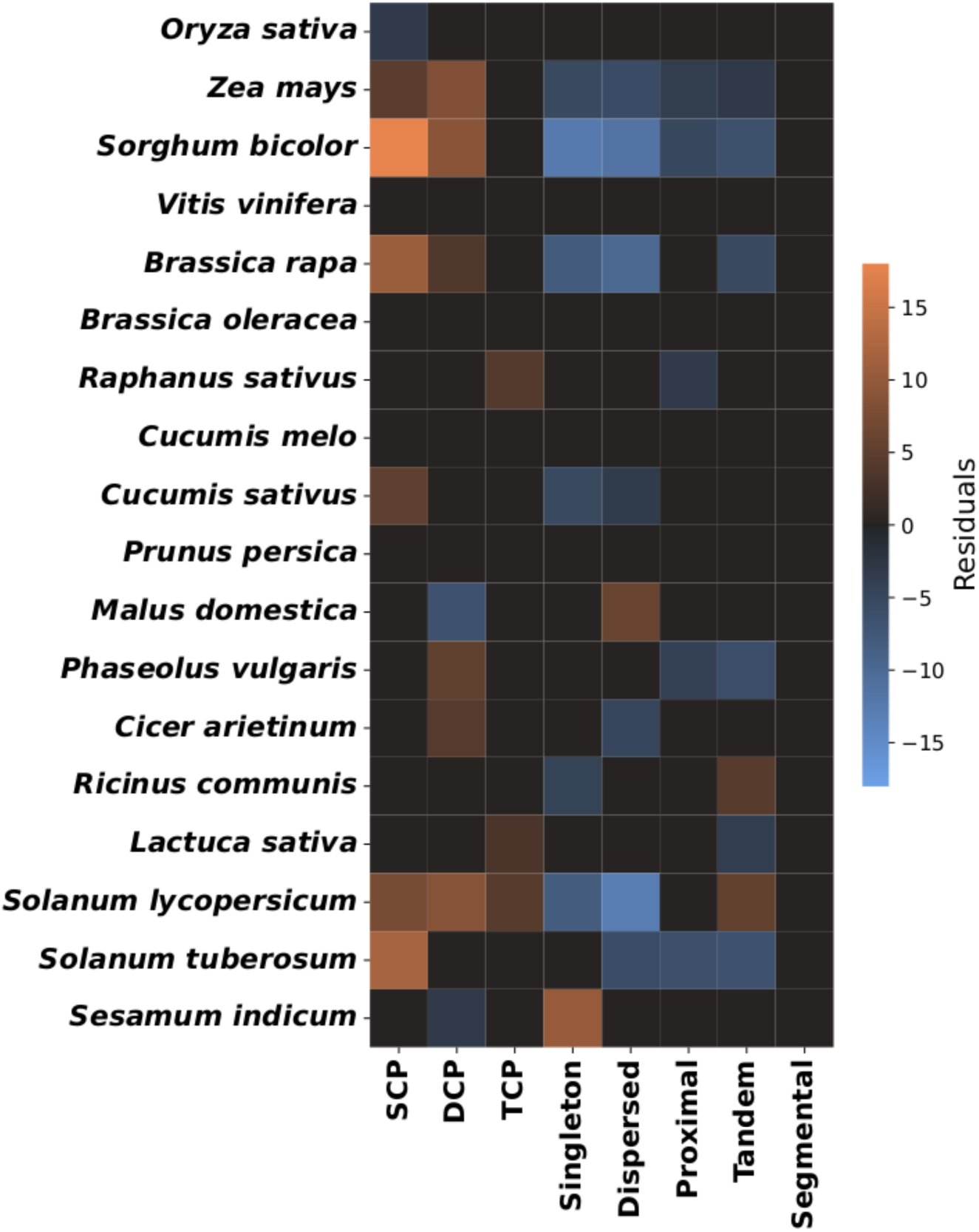
Heat map for the residuals from chi-squared tests comparing the distribution of duplicate-gene classes in candidate domestication genes lists and the genomes of the 18 diploid species tested. Red colors indicate the overrepresentation of particular duplicate-gene classes in the candidate lists following positive chi-square results, blue represents a significant underrepresentation. SCP, DCP, and TCP are acronyms for single-copy, double-copy, and triple-copy paleologs respectively.

**Figure 5:**
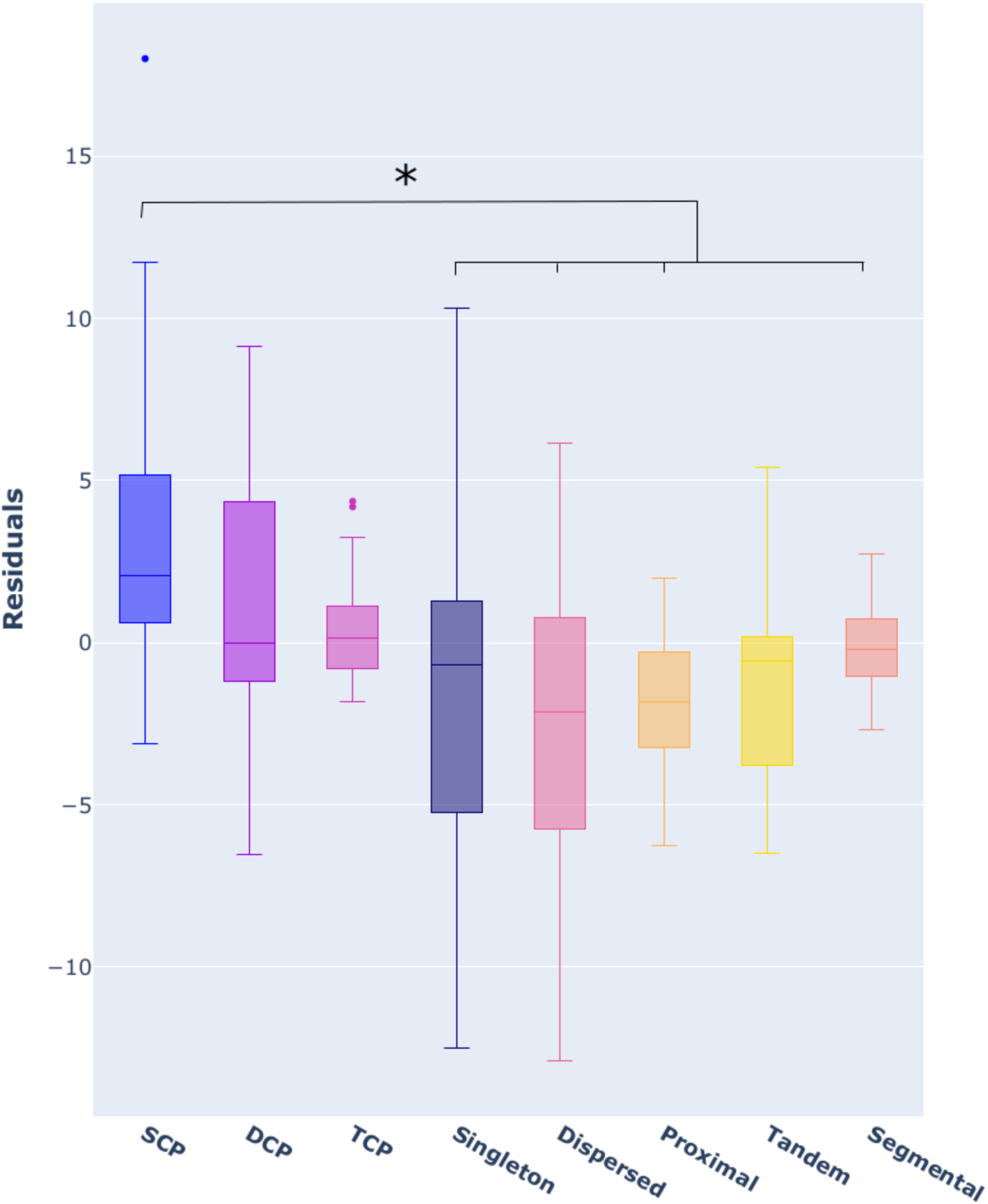
Box and whisker plots comparing the residuals of each duplicate-gene class across all the diploid species tested. The asterisk denotes a positive result from a Friedman test on the ranked residuals of each chi-square test and a Nemenyi post hoc test comparing each gene category in a pairwise fashion.^72,73^

**Figure 6:**
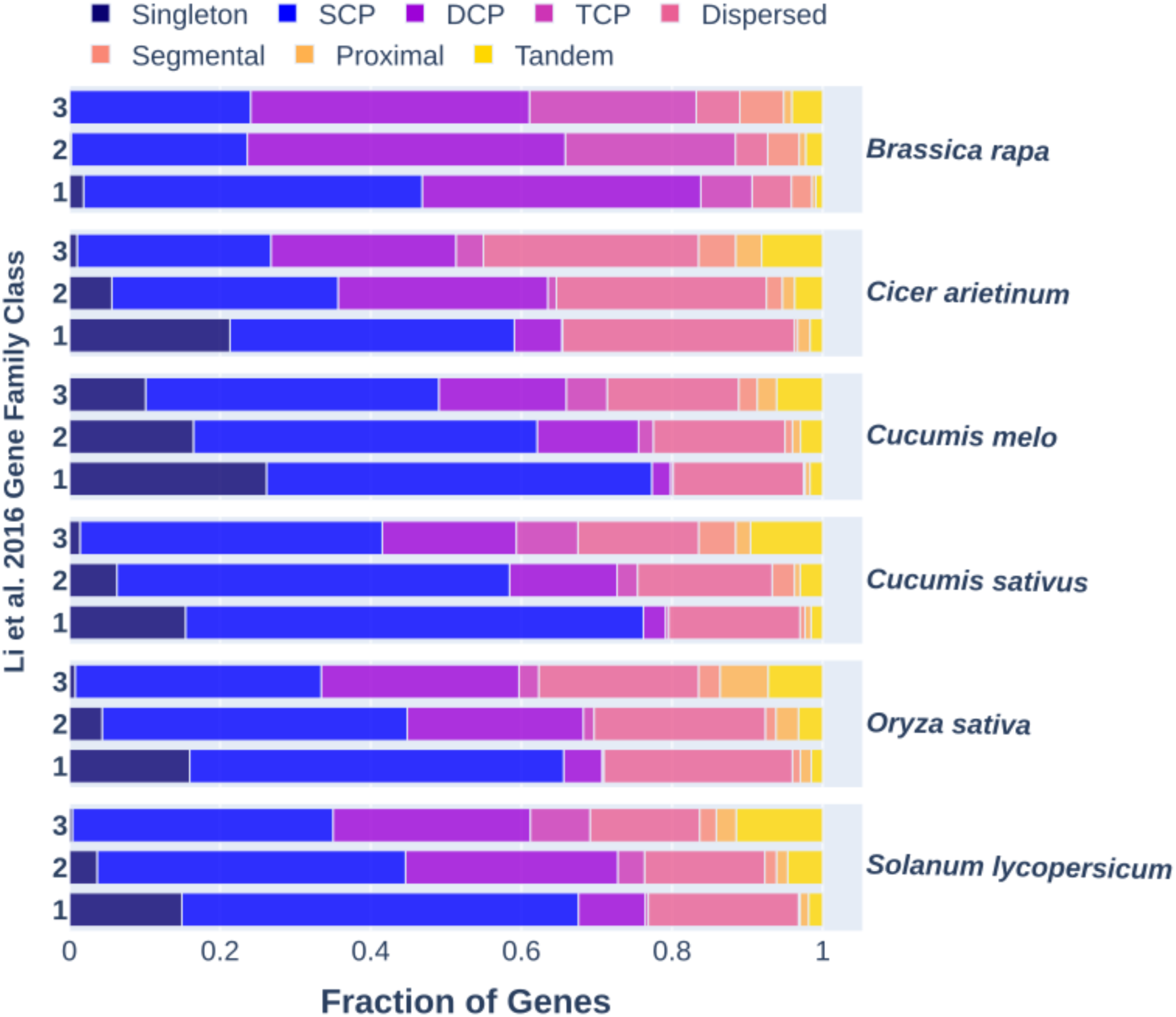
Bar charts showing the fraction of Li et al.^39^ multi-copy (3), intermediate (2), and single-copy (1) gene family classes predicted as paleolog, singletons, and SSD across five species. SCP, DCP, and TCP are acronyms for single-copy, double-copy, and triple-copy paleologs respectively.

### Core-Gene Family Status Influences Domestication Gene Enrichment

Previous work by Li et al.^39^ and Tasdighian et al.^40^ classified gene families found across multiple angiosperm species by their propensity to return to single-copy state following duplication. These studies focussed on gene families shared across the majority of flowering plants they sampled, termed “core-gene families”. To assess whether these duplicability classifications predict involvement in crop domestication, we repeated the chi-squared, post hoc, and Friedman tests using the Li et al.^39^ non-core, single-copy, intermediate, and multicopy core-gene classes in place of the Frackify and MCScanX classifications. There were six species that overlapped in our study.^39,40^ Chi-squared tests were significant across all six species (Supplementary Table 8), indicating that gene family evolutionary history influences their propensity to be involved in domestication. Post-hoc analyses showed that single-copy, intermediate, and multicopy core gene families were significantly enriched in candidate domestication gene lists of three, three, and four species, respectively (Supplementary Table 8). Non-core gene families were significantly depleted from candidate domestication gene lists in five out of the six species sampled (Supplementary Table 8). Despite these patterns, a Friedman test on ranked residuals across duplicability classes was non-significant (X^2^ =6.2, p-value=0.10), suggesting that duplicability *sensu* Li et al.^39^ and Tasdighian et al.^40^ is not a predictor of gene representation in candidate lists; rather, core gene families are more likely to be represented than non-core gene families.

## Discussion

Phylogenetic analyses have shown an association between paleopolyploid ancestry and crop domestication,^1^ but the genomic basis of this association has remained unclear. Previous work in *Brassica rapa* found that paleologs are overrepresented among candidate domestication genes,^22^ raising the question of whether this pattern is specific to *Brassica* or generalizable across angiosperms. Here, we find that paleologs are significantly overrepresented in candidate domestication gene lists in 14 of 22 crop species surveyed, with no other duplicate-gene class showing a comparable pattern of enrichment. Single-copy paleologs—genes that have returned to single-copy status following WGD—were the most consistently enriched class across species. Three species—*O. sativa*, *M. domestica*, and *S. indicum*—showed significant paleolog underrepresentation, variation likely reflecting differences in candidate list size and species-specific histories of post-WGD genome evolution. In contrast, genes derived from small-scale duplications were consistently underrepresented. Despite this variation, paleolog enrichment was independent of both WGD age and the degree of subsequent gene loss, suggesting that the genomic consequences of ancient polyploidy persist as a substrate for crop evolution across tens of millions of years of divergence. All gene duplication classes were present in candidate domestication gene lists across our sample, indicating that SSD and singletons are not precluded from contributing to crop domestication. Nevertheless, the consistent enrichment of paleologs suggests that some feature of genes derived from WGD conferred a selective advantage during domestication of multiple crop species.

Several processes could explain the preferential representation of single-copy paleologs in candidate domestication gene lists. First, selection may be more effective on single-copy paleologs because they are not subjected to the masking effects that limit selection efficacy on multi-copy genes.^47–50^ Second, single-copy paleologs can leverage the elevated genetic diversity that accumulates in paleologs,^22^ variation that becomes more readily available for selection once genes return to single-copy status. Third, purifying selection may favor maintaining at least one functional copy of essential paleologs to avoid yield losses associated with complete gene deletion.^20,21,51,52^ These explanations are not mutually exclusive, and our comparison with gene family duplicability classifications^39,40^ suggests the relationship between dosage constraint and selection is nuanced. Core-gene families of all duplicability statuses—whether rapidly returning to single-copy or persistently retained in duplicate—were enriched in domestication candidates, while duplicability itself was not a significant predictor of representation. This indicates that being a paleolog matters more for domestication potential than the rate at which duplicates are lost. Constraint on copy number does not necessarily preclude selection on allelic variation, and the retention patterns described in prior studies were not direct tests of selection during domestication.^39,40^ We propose direct tests of selection through multi-generation artificial selection experiments or targeted knockdowns of single- and multi-copy paleologs. These tests could assess what aspects of paleologs make them selectively advantageous and if masking is an ongoing phenomenon in paleologs despite having originated millions of years ago.

Regardless of retention status, paleologs were the only gene class to consistently be enriched in candidate gene lists across multiple crop species. In particular, SSD were consistently under-represented in candidate domestication gene lists. This may be counterintuitive as SSD are often observed to be involved in adaptation to abiotic and biotic stress^53–56^ and metabolic diversification.^57^ However the action of artificial selection on genes may be nuanced and determined by the degree of pleiotropy.^32,36,58,59^ Genes central to PPI and GRNs—hubs—are more likely to have increased epistatic and pleiotropic effects than peripheral genes—spokes—resulting in larger phenotypic effects.^32,36,37^ Duplication can decrease pleiotropy and increase the efficacy of positive selection,^31^ but this effect is less pronounced in spokes relative to hubs. Hub genes are also less likely to be retained following SSD due to their negative effects on dosage balance.^14,26,29,37^ Under this hypothesis, paleologs, or the gene families they are enriched within, would be favored by artificial selection over SSD due to their larger effect on pleiotropy, increased phenotypic effects, and long term impacts on GRN motif evolution.^37^ This hub-targeting effect may contribute to convergent domestication syndromes across species, as artificial selection repeatedly acts on similar network control points. Similar to our discussion of copy-number variation, it is important to emphasize these effects do not preclude SSD from being selected upon during domestication. SSD-derived gene classes were present in candidate lists of each species, and over represented in three crops. However, there is a clear bias in representation towards paleologs and against SSD that requires additional research to mechanistically explain. Future network analyses could test whether paleologs occupy systematically different positions in protein-protein interaction networks than SSD, which would help distinguish whether network centrality or duplication history drives the observed patterns.

There are several caveats when interpreting results from candidate gene lists due to the nature of their construction. Many methods for identifying candidate gene lists, such as scans for selection, report the target of selection along with neighboring genes. These scans are also known to be biased due to recombination rate heterogeneity and genome structural variation^60–62^, which are also correlated with rates of gene deletion and the deterioration of syntenic blocks.^63–65^ Given conserved synteny was used for paleolog detection^42^, it is possible there is a bias in how paleologs and non-paleologs are detected during scans for selection. The enrichment of multi-copy paleologs in sweep-based analyses could reflect physical linkage to selected single-copy paleologs rather than direct selection on multi-copy genes—a hypothesis testable through detailed linkage disequilibrium analysis. Similarly, tests of selection that target specific loci, such as the McDonald-Kreitman test, decrease in power with lower sequence divergence and polymorphism.^66,67^ Given SSD are often younger than paleologs^14,17,68^ and show lower genetic diversity^22^, it is plausible they are less likely to be seen by tests like McDonald-Kreitman. Although biases in how candidate lists are made could plausibly favor paleologs over other gene duplication classes, they likely cannot fully explain the consistent over representation of paleologs in candidate lists. Across studies of the 14 species exhibiting paleolog enrichment, six methods were used to generate candidate lists (Table S1). These methods include tests such as Genome Wide Association Studies (GWAS) that have no known biases towards paralogs.

Here we show that paleologs were frequently selected for during the domestication of crops, with a bias towards those that returned to single-copy state. Although other duplicate-gene classes were involved in domestication of multiple crop species, there is a bias against SSD in candidate domestication gene lists. Popular models of duplicate gene evolution that invoke gene-functional categories and dosage sensitivity^19,28–30,44–46^ were able to explain the distribution of duplicate-gene classes across our sampled species. However, these models could not fully explain why particular duplicate-gene classes were over- or underrepresented in candidate domestication gene lists. There are several alternative hypotheses for why paleologs were important during domestication, the two most plausible being increased novelty found in paleologs and the masking of deleterious gene deletions on yield. Future work to understand how selection acts on paralogs formed by different modes of duplication and why paleologs were important during domestication will likely give us greater insight into duplicate gene evolution. Given the ubiquity of polyploidy among angiosperms,^10,11,69^ a deeper understanding of how paleopolyploidy continues to impact crops is important for future crop improvement efforts.

## Methods

### Candidate gene list selection

We used Google Scholar to conduct a literature search for publications on candidate domestication genes. Several searches were conducted using combinations of the terms candidate, gene, list, domestication, crops, plants, traits, and agriculture. Publications within the first 10 pages of each query were utilized if they referenced a publicly available reference genome that matched candidate gene list names. The purpose of this filtering step was to ensure gene names for all CDS sequences could be matched between paleolog inferences and candidate lists. Secondary filtering took place to select studies that utilized methodologies that detected direct evidence of selection. Such evidence included reduction of diversity in the form of π, and Fst compared between domestic and landrace species, and studies associating known domesticated phenotypes with candidate genes such as QTL and GWAS Studies (table 1). For several species there were multiple gene lists available that matched to different reference genomes. To maximize our sample size we selected studies that reported lists which contained the largest number of candidates.

### Partitioning genomes into paleologs and non-paleologs

We used the Frackify pipeline to identify paleologs from the most recent WGD in each crop.^42^ The Frackify pipeline requires syntenic and synonymous divergence (K*_S_*) information from the target species and a closely related outgroup that does not share the focal WGD.^42^ For these inferences we utilized MCScanX with default settings.^43^ The K*_a_* and K*_S_* values for each collinear-gene pair were calculated using the add_ka_and_ks_to_collinearity.pl script provided in MCScanX.^43^ We then fit a density curve to the distribution of pairwise collinear gene synonymous divergence (K*_S_*) values from the MCScanX results using the nparam_density and gaussian_kde functions from the Numpy and Scipy python libraries.^70,71^ The maxima of each WGD and orthology peak in the distribution of K*_S_* values was determined using the find_peaks function from the Scipy python library.^71^ Our study included four polyploids, *B. juncea, B. napus, G. hirsutum,* and *G. max.* In polyploids, paleolog copy variation is doubled, with some additional copies being lost over time. Frackify was developed for use in diploids, and any additional copies of each paleologs would result in the misclassification of the retention status of each paleolog. To avoid inaccurate retention inferences, an altered version of Frackify was used that does not determine the retention status of each paleolog. Further information can be found on the Frackify git repo (https://gitlab.com/barker-lab/frackify)

### Small scale duplicate gene classification

In addition to identifying duplicate genes that originate from large scale whole genome duplications, we also identified duplicates originating from more recent small scale duplications. Genes not predicted as paleologs were classified as dispersed, proximal, tandem, or segmental using the duplicate_gene_classifier available in MCScanX.^43^

### Duplicate gene representation in candidate gene lists

We used a Pearson chi-square tests of goodness of fit in each species to compare the distribution of duplicate-gene classes within candidate gene lists to that of the reference genome^72,73^ and a Benjamini and Hochberg multiple test correction from the p.adjust() function available in R.^72,74^ To further assess which duplicate-gene classes contributed the most to model significance we use the chisq.posthoc.test available in R on the residuals of each chi-square test across our species.^72,75^ We use a Friedman test on the ranked residuals of the chi-square tests to test if particular duplicate-gene classes consistently contributed to model significance across all our sampled species.^72,73^ Following the Friedman test we conducted a Nemenyi post hoc test in a pairwise fashion to determine which duplicate-gene classes significantly differed from each other in their ranked residuals.^72,73^

### Phylogenetic analysis

Patterns of gene loss and time since duplication may influence the propensity of duplicate genes to be involved in crop domestication. We assessed how these processes correlate using PGLS to account for the non-phylogenetic independence among species. To begin we accessed a time calibrated phylogeny with 32,223 Angiosperm species and pruned it to include only our focal species.^76,77^ *M. sieversii*, the direct ancestor to *M. domestica,* was used as a substitute tip to represent *M. domestica.^78^* PGLS were performed using the pgls function available in the caper V1.0.1 R.^79^ Kappa and delta were held constant, while lambda was determined by searching the maximum likelihood space between zero and one. For each paleolog retention class, we tested how the fraction of the genome they comprised and divergence time since their duplication correlated with the residuals of each paleolog class from chi-squared test across our species. Additionally, we included the length of candidate gene lists in these tests to account for their variability between species. As Frackify can only make retention status inferences in diploid species, we did not include any polyploid species in these analyses.

### Comparison of gene class inferences to a previous study

Duplicate-gene class inferences in our study may be biased by quality of the reference genome and outgroup genome used. Previous work by Li et al.^39^ assessed core gene families among Angiosperms for their propensity to be lost following duplication and sorted them into three classes. Class one consisted of highly duplication resistant gene families, whereas class three was highly duplicated gene families, and class two was an intermediate class.^39^ To assess how our duplicate-gene class inferences compared to those of Li et al.^39^ we compared the overlap of each gene list for six of the species in our analyses that shared a reference genome with the Li et al.^39^ study. To further assess if propensity of gene families to be lost following duplication impacts their involvement in the domestication process, we repeated the chi-square and post hoc tests using the Li et al.^39^ non-core, core class one, two, and three gene family classifications rather than our duplicate-gene classifications. Following these tests we conducted a Friedman test on the ranked residuals of the chi-squared tests in each species.

### GO Term Enrichment

To better understand the functions duplicate genes represent we used GoGetter to functionally annotate each reference genome we surveyed.^80^ Annotations were made using the TAIR10 GOSlim database.^81,82^ For each duplicate-gene class and our candidate gene lists we tested the under or over representation of each GO category by comparing against the fraction represented in the reference genome using a chi-square test and assessed the contribution of each duplicate-gene class to model significance using the chisq.posthoc.test available in R.^72,75^ To test if particular GO categories were consistently over or underrepresented in each duplicate-gene class across all our sampled species we conducted a Friedman test and a Nemenyi post hoc test on the ranked residuals of each GO category from the chi-squared tests

## Data Availability

MCScanX, Frackify, and GO term results are available at https://gitlab.com/barker-lab/McKibben_Barker_2026_Paleolog_Domestication. All other data are available in the Article and Supplementary Information.

## Code Availability

Frackify is currently available at https://gitlab.com/barker-lab/frackify. The scripts used for statistical analyses conducted in this manuscript are available at https://gitlab.com/barker-lab/McKibben_Barker_2026_Paleolog_Domestication.

## Acknowledgements

We would like to thank Dr. Justin Conover from the Danforth Plant Science Center, Dr. Geoffery Finch from the University of Arizona, and Dr. Sylvia Kinosian from Montana State University for their comments and discussion of this manuscript. This material is based upon High Performance Computing (HPC) resources supported by the University of Arizona TRIF, UITS, and Research, Innovation, and Impact (RII) and maintained by the UArizona Research Technologies department. M.S.B. is supported by the CAMBIUM NRT under NSF Grant No. 2346054.

## Supplementary Information

**Supplementary Table 1:** Table summarizing the species, genomes, and candidate gene lists utilized in this study. Column one is the latin species name, two the method or metric used to identify candidate genes, three the general traits being selected for in the associated study, and five the number of candidate genes. Columns six through eight summarize the reference genome and candidate lists sources. Columns nine and ten summarize the outgroup used for the Frackify analysis and their source.

**Supplementary Table 2:** Table summarizing the number of each gene category in each diploid species and gene lists. Column one states the species name and column two states the gene lists associated with the appropriate row; either the genome or candidate gene lists for the associated species. Column three is the median Ks value for the focal WGD peak in the Ks plot being analyzed by Frackify. The remaining columns represent the number of each duplicate-gene class in the associated gene lists. SCP, DCP, and TCP are acronyms for single-copy, double-copy, and triple-copy paleologs.

**Supplementary Table 3:** Table summarizing the number of each gene category in each polyploid species and gene lists. Column one states the species name and column two states the gene lists associated with the appropriate row; either the genome or candidate gene lists for the associated species. Column three is the median Ks value for the focal WGD peak in the Ks plot being analyzed by Frackify. The remaining columns represent the number of each duplicate-gene class in the associated gene lists. SCP, DCP, and TCP are acronyms for single-copy, double-copy, and triple-copy paleologs.

**Supplementary Table 4:** Chi-squared results for each diploid species. Column one states the species being tested, two the X^2^ statistic, three the degrees of freedom, and four the p-value from the test. The remaining columns are the raw residuals from the chi-squared test for each duplicate-gene class and the associated p-value from the post hoc tests on each gene category. SCP, DCP, and TCP are acronyms for single-copy, double-copy, and triple-copy paleologs.

**Supplementary Table 5:** Chi-squared results for each polyploid species. Column one states the species being tested, two the X^2^ statistic, three the degrees of freedom, and four the p-value from the test. The remaining columns are the raw residuals from the chi-squared test for each duplicate-gene class and the associated p-value from the post hoc tests on each gene category. SCP, DCP, and TCP are acronyms for single-copy, double-copy, and triple-copy paleologs.

**Supplementary Table 6:** Table for the p-values of pairwise comparisons of each duplicate-gene class across all diploid species tested from a Nemenyi post-hoc test following a Friedman test.

**Supplementary Table 7:** The distribution of duplicate-gene classes predicted by Frackify and MCScanX in each gene family class from Li et al.^39^. Gene family class 1 are genes that are retained primarily in single-copy state across sampled angiosperms, class 2 are an intermediate class, and 3 are gene families primarily retained in duplicate. SCP, DCP, and TCP are acronyms for single-copy, double-copy, and triple-copy paleologs. Columns one and two denote the associated species tested and the duplicate-gene class being analyzed respectively.

**Supplementary Table 8:** Chi-squared results comparing the enrichment of each gene family class from Li et al.^39^ in the candidate gene lists compared to the reference genome. Column one states the species being tested, two the X^2^ statistic, three the degrees of freedom, and four the p-value from the test. The remaining columns are the raw residuals from the chi-squared test for each duplicability class and the associated p-value from the post hoc tests on each gene category. Gene family class 1 are genes that are retained primarily in single-copy state across sampled angiosperms, class 2 are an intermediate class, and 3 are gene families primarily retained in duplicate.

**Supplementary Table 9:** Chi-squared results comparing the distribution of GO functional categories in each duplicate-gene class compared to the reference genome. Column one is the species, two the duplicate-gene class within the associated species being tested, three the X^2^ statistic, and four the p-value from the test.

**Supplementary Tables 10-13**: Tables for the p-values of pairwise comparisons of each GO functional category across all species using a Nemenyi post-hoc test following a Friedman test. Table 10 are the results of the Nemenyi post-hoc test in double-copy paleologs, Table 11 single-copy paleologs, Table 12 SSD, and Table 13 the candidate domestication gene lists in each species.

## Notes

https://gitlab.com/barker-lab/McKibben_Barker_2026_Paleolog_Domestication

### Competing Interest Statement

The authors have declared no competing interest.

### Summary of Updates

We have added additional clarity to the abstract and discussion sections. We also corrected minor typos and inconsistencies throughout the manuscript. Finally, we edited an address for the author affiliations.

## References

1. Salman-Minkov, A., Sabath, N. & Mayrose, I. Whole-genome duplication as a key factor in crop domestication. Nat Plants 2, 16115 (2016).

2. Akagi, T., Jung, K., Masuda, K. & Shimizu, K. K. Polyploidy before and after domestication of crop species. Curr. Opin. Plant Biol. 69, 102255 (2022).

3. Paterson, A. H. Polyploidy, evolutionary opportunity, and crop adaptation. Genetics of Adaptation 191–196 (2005).

4. Zhang, K., Wang, X. & Cheng, F. Plant Polyploidy: Origin, evolution, and its influence on crop domestication. Horticultural Plant Journal 5, 231–239 (2019).

5. Pecinka, A., Fang, W., Rehmsmeier, M., Levy, A. A. & Mittelsten Scheid, O. Polyploidization increases meiotic recombination frequency in *Arabidopsis*. BMC Biology 9, 1–7 (2011).

6. Renny-Byfield, S. & Wendel, J. F. Doubling down on genomes: polyploidy and crop plants. Am. J. Bot. 101, 1711–1725 (2014).

7. Selmecki, A. M. et al. Polyploidy can drive rapid adaptation in yeast. Nature 519, 349–352 (2015).

8. Baniaga, A. E., Marx, H. E., Arrigo, N. & Barker, M. S. Polyploid plants have faster rates of multivariate niche differentiation than their diploid relatives. Ecology Letters 23, 68–78 (2020).

9. Van de Peer, Y., Ashman, T.-L., Soltis, P. S. & Soltis, D. E. Polyploidy: an evolutionary and ecological force in stressful times. Plant Cell 33, 11–26 (2021).

10. 1KP Transcriptomes Initiative. One thousand plant transcriptomes and the phylogenomics of green plants. Nature 574, 679–685 (2019).

11. McKibben, M. T. W., Finch, G. & Barker, M. S. Species-tree topology impacts the inference of ancient whole-genome duplications across the angiosperm phylogeny. American Journal of Botany 111, e16378 (2024).

12. Van de Peer, Y., Maere, S. & Meyer, A. The evolutionary significance of ancient genome duplications. Nature Reviews Genetics 10, 725–732 (2009).

13. Li, Z. et al. Patterns and processes of diploidization in land plants. Annu. Rev. Plant Biol. 72, 387–410 (2021).

14. Lynch, M. & Conery, J. S. The evolutionary fate and consequences of duplicate genes. Science 290, 1151–1155 (2000).

15. Lynch, M. & Conery, J. S. The evolutionary demography of duplicate genes. Genome Evolution 35–44 (2003).

16 Ohno, S. The creation of a new gene from a redundant duplicate of an old gene. in Evolution by Gene Duplication (ed. Ohno, S.) 71–82 (Springer Berlin Heidelberg, Berlin, Heidelberg, 1970).

17. Hakes, L., Pinney, J. W., Lovell, S. C., Oliver, S. G. & Robertson, D. L. All duplicates are not equal: the difference between small-scale and genome duplication. Genome Biol. 8, 1–13 (2007).

18. Chapman, B. A., Bowers, J. E., Feltus, F. A. & Paterson, A. H. Buffering of crucial functions by paleologous duplicated genes may contribute cyclicality to angiosperm genome duplication. Proc. Natl. Acad. Sci. U. S. A. 103, 2730–2735 (2006).

19. Conant, G. C., Birchler, J. A. & Pires, J. C. Dosage, duplication, and diploidization: clarifying the interplay of multiple models for duplicate gene evolution over time. Curr. Opin. Plant Biol. 19, 91–98 (2014).

20. Huang, K. et al. The genomics of linkage drag in inbred lines of sunflower. Proc. Natl. Acad. Sci. U. S. A. 120, e2205783119 (2023).

21. Lee, J. S. et al. Expression complementation of gene presence/absence polymorphisms in hybrids contributes importantly to heterosis in sunflower. J. Adv. Res. 42, 83–98 (2022).

22. Qi, X. et al. Genes derived from ancient polyploidy have higher genetic diversity and are associated with domestication in *Brassica rapa*. New Phytol. 230, 372–386 (2021).

23. Freeling, M., Scanlon, M. J. & Fowler, J. E. Fractionation and subfunctionalization following genome duplications: mechanisms that drive gene content and their consequences. Curr. Opin. Genet. Dev. 35, 110–118 (2015).

24. Walsh, B. Population-genetic models of the fates of duplicate genes. Genetica 118, 279–294 (2003).

25. Scannell, D. R. & Wolfe, K. H. A burst of protein sequence evolution and a prolonged period of asymmetric evolution follow gene duplication in yeast. Genome Res. 18, 137–147 (2008).

26. Hahn, M. W. Distinguishing among evolutionary models for the maintenance of gene duplicates. J. Hered. 100, 605–617 (2009).

27. Des Marais, D. L. & Rausher, M. D. Escape from adaptive conflict after duplication in an anthocyanin pathway gene. Nature 454, 762–765 (2008).

28. Vance, Z. & McLysaght, A. Ohnologs and SSD Paralogs Differ in Genomic and Expression Features Related to Dosage Constraints. Genome Biol. Evol. 15, (2023).

29. Edger, P. P. & Pires, J. C. Gene and genome duplications: the impact of dosage-sensitivity on the fate of nuclear genes. Chromosome Res. 17, 699–717 (2009).

30. Freeling, M. Bias in plant gene content following different sorts of duplication: tandem, whole-genome, segmental, or by transposition. Annu. Rev. Plant Biol. 60, 433–453 (2009).

31. Nocchi, G., Whiting, J. R. & Yeaman, S. Repeated global adaptation across plant species. Proc Natl Acad Sci U S A 121, e2406832121 (2024).

32. Whiting, J. R. et al. The genetic architecture of repeated local adaptation to climate in distantly related plants. Nat Ecol Evol 8, 1933–1947 (2024).

33. Gasparini, K., Moreira, J. D. R., Peres, L. E. P. & Zsögön, A. De novo domestication of wild species to create crops with increased resilience and nutritional value. Curr Opin Plant Biol 60, 102006 (2021).

34. Meyer, R. S., DuVal, A. E. & Jensen, H. R. Patterns and processes in crop domestication: an historical review and quantitative analysis of 203 global food crops. New Phytol. 196, 29–48 (2012).

35. Hammer, K. Доместикационный синдром. Kulturpflanze 32, 11–34 (1984).

36. Pouzet, S. & Le Rouzic, A. Gene network topology drives the mutational landscape of gene expression. bioRxiv 2024.11.28.625874 (2024) doi:10.1101/2024.11.28.625874.

37. Almeida-Silva, F. & Van de Peer, Y. Whole-genome Duplications and the Long-term Evolution of Gene Regulatory Networks in Angiosperms. Mol. Biol. Evol. 40, (2023).

38. Wagner, A. Evolution of gene networks by gene duplications: a mathematical model and its implications on genome organization. Proceedings of the National Academy of Sciences 91, 4387–4391 (1994).

39. Li, Z., et al. Gene duplicability of core genes is highly consistent across all angiosperms. The Plant Cell vol. 28 326–344 Preprint at 10.1105/tpc.15.00877 (2016).

40. Tasdighian, S. et al. Reciprocally Retained Genes in the Angiosperm Lineage Show the Hallmarks of Dosage Balance Sensitivity. Plant Cell 29, 2766–2785 (2017).

41. Carretero-Paulet, L. & Van de Peer, Y. The evolutionary conundrum of whole-genome duplication. Am J Bot 107, 1101–1105 (2020).

42. McKibben, M. T. W. & Barker, M. S. Applying Machine Learning to Classify the Origins of Gene Duplications. in Polyploidy: Methods and Protocols (ed. Van de Peer, Y.) 91–119 (Springer US, New York, NY, 2023).

43. Wang, Y. et al. MCScanX: a toolkit for detection and evolutionary analysis of gene synteny and collinearity. Nucleic Acids Res. 40, e49 (2012).

44. Wang, Y. et al. Modes of gene duplication contribute differently to genetic novelty and redundancy, but show parallels across divergent angiosperms. PLoS One 6, e28150 (2011).

45. Birchler, J. A., Riddle, N. C., Auger, D. L. & Veitia, R. A. Dosage balance in gene regulation: biological implications. Trends Genet. 21, 219–226 (2005).

46. Freeling, M. & Thomas, B. C. Gene-balanced duplications, like tetraploidy, provide predictable drive to increase morphological complexity. Genome Res. 16, 805–814 (2006).

47. Tanaka, K. M., Takahasi, K. R. & Takano-Shimizu, T. Enhanced fixation and preservation of a newly arisen duplicate gene by masking deleterious loss-of-function mutations. Genet Res (Camb) 91, 267–280 (2009).

48. Gossmann, T. I. & Schmid, K. J. Selection-driven divergence after gene duplication in *Arabidopsis thaliana*. J Mol Evol 73, 153–165 (2011).

49. Lynch, M., O’Hely, M., Walsh, B. & Force, A. The probability of preservation of a newly arisen gene duplicate. Genetics 159, 1789–1804 (2001).

50. Lynch, M. The evolution of genetic networks by non-adaptive processes. Nat Rev Genet 8, 803–813 (2007).

51. Dopman, E. B. & Hartl, D. L. A portrait of copy-number polymorphism in *Drosophila melanogaster*. Proc Natl Acad Sci U S A 104, 19920–19925 (2007).

52. Katju, V. & Bergthorsson, U. Copy-number changes in evolution: rates, fitness effects and adaptive significance. Front Genet 4, 273 (2013).

53. Du, L., Ma, Z. & Mao, H. Duplicate genes contribute to variability in abiotic stress resistance in allopolyploid wheat. Plants 12, 2465 (2023).

54. Zhang, Y. et al. Expression partitioning of homeologs and tandem duplications contribute to salt tolerance in wheat (Triticum aestivum L.). Sci Rep 6, 21476 (2016).

55. Dossa, K., Diouf, D. & Cissé, N. Genome-wide investigation of hsf genes in sesame reveals their segmental duplication expansion and their active role in drought stress response. Front Plant Sci 7, 1522 (2016).

56. Wu, H.-J. et al. Insights into salt tolerance from the genome of *Thellungiella salsuginea*. Proc Natl Acad Sci U S A 109, 12219–12224 (2012).

57. Bird et al. Phylogenetic and genomic mechanisms shaping glucosinolate innovation. Current Opinion in Plant Biology 85, 102705 (2025).

58. Wang, Z., Liao, B.-Y. & Zhang, J. Genomic patterns of pleiotropy and the evolution of complexity. Proceedings of the National Academy of Sciences 107, 18034–18039 (2010).

59. Wagner, G. P. et al. Pleiotropic scaling of gene effects and the ‘cost of complexity’. Nature 452, 470–472 (2008).

60. Lotterhos, K. E. The effect of neutral recombination variation on genome scans for selection. G3 9, 1851–1867 (2019).

61. Matthey-Doret, R. & Whitlock, M. C. Background selection and F : Consequences for detecting local adaptation. Mol. Ecol. 28, 3902–3914 (2019).

62. Booker, T. R., Yeaman, S. & Whitlock, M. C. Variation in recombination rate affects detection of outliers in genome scans under neutrality. Mol. Ecol. 29, 4274–4279 (2020).

63. Ferguson, S., Jones, A., Murray, K., Schwessinger, B. & Borevitz, J. O. Interspecies genome divergence is predominantly due to frequent small scale rearrangements in Eucalyptus. Mol. Ecol. 32, 1271–1287 (2023).

64. Akhunov, E. D. et al. Synteny perturbations between wheat homoeologous chromosomes caused by locus duplications and deletions correlate with recombination rates. Proc. Natl. Acad. Sci. U. S. A. 100, 10836–10841 (2003).

65. Gaut, B. S., Wright, S. I., Rizzon, C., Dvorak, J. & Anderson, L. K. Recombination: an underappreciated factor in the evolution of plant genomes. Nat. Rev. Genet. 8, 77–84 (2007).

66. Zhai, W., Nielsen, R. & Slatkin, M. An investigation of the statistical power of neutrality tests based on comparative and population genetic data. Mol. Biol. Evol. 26, 273–283 (2009).

67. Parsch, J., Zhang, Z. & Baines, J. F. The influence of demography and weak selection on the McDonald-Kreitman test: an empirical study in Drosophila. Mol. Biol. Evol. 26, 691–698 (2009).

68. Rody, H. V. S., Baute, G. J., Rieseberg, L. H. & Oliveira, L. O. Both mechanism and age of duplications contribute to biased gene retention patterns in plants. BMC Genomics 18, 46 (2017).

69. Landis, J. B. et al. Impact of whole-genome duplication events on diversification rates in angiosperms. Am. J. Bot. 105, 348–363 (2018).

70. Harris, C. R. et al. Array programming with NumPy. Nature 585, 357–362 (2020).

71. Virtanen, P. et al. SciPy 1.0: fundamental algorithms for scientific computing in Python. Nat. Methods 17, 261–272 (2020).

72. Team, R. C. R Core Team-RA Language and Environment for Statistical Computing. R Foundation for Statistical Computing, Vienna, Austria. (2022).

73. Hollander, M., Wolfe, D. A. & Chicken, E. Nonparametric Statistical Methods. (John Wiley & Sons, 2013).

74. Benjamini, Y. & Hochberg, Y. Controlling the false discovery rate: a practical and powerful approach to multiple testing. J. R. Stat. Soc. Ser. B. Stat. Methodol. 57, 289–300 (2018).

75. Mark Beasley, T. & Schumacker, R. E. MULTIPLE REGRESSION APPROACH TO ANALYZING CONTINGENCY TABLES: POST HOC AND PLANNED COMPARISON PROCedures. The Journal of Experimental Education 79–93 (2010).

76. Zanne, A. E. et al. Corrigendum: Three keys to the radiation of angiosperms into freezing environments. Nature 521, 380 (2015).

77. Paradis, E., Claude, J. & Strimmer, K. APE: Analyses of phylogenetics and evolution in r language. Bioinformatics 20, 289–290 (2004).

78. Velasco, R. et al. The genome of the domesticated apple (*Malus× domestica Borkh*.). Nat. Genet. 42, 833–839 (2010).

79. Orme, D. et al. The caper package: comparative analysis of phylogenetics and evolution in R. R package version 5, 1–36 (2013).

80. Sessa, E. B., Masalia, R. R., Arrigo, N., Barker, M. S. & Pelosi, J. A. GOgetter: A pipeline for summarizing and visualizing GO slim annotations for plant genetic data. Appl. Plant Sci. 11, e11536 (2023).

81. Berardini, T. Z. et al. The *Arabidopsis* information resource: Making and mining the ‘gold standard’ annotated reference plant genome. Genesis 53, 474–485 (2015).

82. Berardini, T. Z. et al. Functional Annotation of the *Arabidopsis* Genome Using Controlled Vocabularies. Plant Physiol. 135, 745–755 (2004).

